# Similarity in evoked responses does not imply similarity in macroscopic network states across tasks

**DOI:** 10.1101/2021.11.27.470015

**Authors:** Javier Rasero, Richard Betzel, Amy Isabella Sentis, Thomas E. Kraynak, Peter J. Gianaros, Timothy Verstynen

**Affiliations:** Department of Psychology, Carnegie Mellon University, Pittsburgh, PA 15213, USA; Neuroscience Institute, Carnegie Mellon University, Pittsburgh, PA 15213, USA; Department of Psychological and Brain Sciences, Indiana University, Bloomington, IN 47405, USA; Center for the Neural Basis of Cognition, University of Pittsburgh and Carnegie Mellon University, Pittsburgh, PA 15213, USA; Department of Psychology, University of Pittsburgh, Pittsburgh, PA 15260, USA; Biomedical Engineering, Carnegie Mellon University, Pittsburgh, PA 15213, USA

## Abstract

There is an ongoing debate as to whether cognitive processes arise from a group of functionally specialized brain modules (modularism) or as the result of distributed nonlinear processes (dynamical systems theory). The former predicts that tasks recruiting similar local brain areas should be equally similar in their network profiles. The latter allows for differential connectivity, even when the areas recruited are largely the same. Here we evaluated both views at the macroscopic level by comparing region-wise activation patterns and functional correlation profiles from a large sample of healthy subjects (N=242) that performed two executive control tasks known to recruit nearly identical brain areas, the color-word Stroop task and the Multi-Source Interference Task (MSIT). Using a measure of instantaneous functional correlations, based on edge time series, we estimated the task-related networks that differed between incongruent and congruent conditions. At the group level, the two tasks were much more different in their network profiles than in their evoked activity patterns. This is found even when matching the degrees of freedom of both activation patterns and functional correlation profiles, when considering subject-level differences, after changing brain parcellations, and if employing alternative methods for defining task-related network profiles. Our results are consistent with the perspective of the brain as a dynamical system, suggesting that task representations should be independently evaluated at both node and edge (connectivity) levels.

**Significant Statement:** If the brain is strictly modular at the macroscopic scale, then recruiting the same brain regions should result in the same functional interactions between regions. However, if the brain is a dynamical system, with information represented at both the node and edge levels, then two tasks could have the same pattern of activation, but largely different functional correlation profiles. Here we tested this contrastive prediction using two tasks with overlapping cognitive demands, but different sensory signals. Despite being nearly identical in their activation patterns, we found that the tasks produced largely different functional correlation profiles. These findings reinforce the view of the brain as a dynamical system, with task states represented both within and across regions.

## Introduction

The idea of a modular mind (Fodor, 1983), where cognition arises from the interplay between specialized, domain-specific units that represent fundamental cognitive processes, has dominated the cognitive neuroscientific view of the brain since the its inception (e.g., Posner et al., 1988). Here the cognitive “modules” are mapped to unique brain areas that execute specific processes (e.g., detecting specifics sound frequencies, estimating value, contracting specific muscle groups) (Feinberg and Farah, 2006). Over the last three decades, this modular view of the brain has largely been justified from empirical observations using non-invasive brain imaging methods, like positron emission tomography and functional MRI (fMRI), where experiments and analytical methods were explicitly designed to isolate clusters of regions aligned to certain functional domains, such as vision (e.g., Bihan et al., 1993), control (e.g., Porro et al., 1996), language (e.g., Binder et al., 1997), or affect (e.g., Anders et al., 2004). A critical assumption of this modularist perspective is that task-relevant representations are strictly limited to the domain-specific modules (e.g., specific brain regions), with communication between modules simply being a matter of relaying information from one stage of processing to the next. In other words, co-activating the same two modules (i.e., brain regions) in the same way across two different tasks will also lead to similar connectivity profiles between those modules.

This picture of a modular brain has been progressively challenged over the years by the idea that cognition arises from a dynamical system. From this perspective, a given cognitive function cannot be solely understood by investigating its components separately, but requires also understanding the interactions between units as well (Gelder, 1995). While almost as old as the modularist view of the brain, this dynamical systems perspective has gained ground over the past decade in systems neuroscience, where multi-unit recording studies have shown that task representations emerge as a low-dimensional manifold of population activity, both within and between brain areas (Russo et al., 2020; Churchland et al., 2012; Sadtler et al., 2014; Oby et al., 2019). This observation at the microscale level extends to observations of macroscopic brain dynamics as well (e.g., Kriegeskorte et al., 2008; Ejaz et al., 2015). Indeed, with the rise of connectomics (Behrens and Sporns, 2012), the idea of the brain as a dynamical network (Sporns, 2013), where information is also encoded *between* units (Crossley et al., 2013; Yeo et al., 2014; Bertolero et al., 2015), has gained popularity in cognitive neuroscience. A contrasting assumption of the dynamical systems view, compared to the modularist view, is that task-relevant representations are encoded by both the individual regions and the communication dynamics between those regions. Therefore, two tasks may activate the same pattern of nodes, but express dissimilar network profiles at the edges (Prinz et al., 2004; Hooper, 2004).

Here we sought to shed light on this debate by testing whether the apparent overlapping patterns of blood oxygen level-dependent (BOLD) activation that are elicited during two response conflict tasks (Sheu et al., 2012), the color-word Stroop task (Stroop, 1935) and the Multi-Source Interference Task (MSIT) (Bush and Shin, 2006), also have similar task-related network profiles. Our study develops on previous work exploring the relationship between task activation and functional correlations (Alnæs et al., 2015; Chan et al., 2017; Gratton et al., 2016; Krienen et al., 2014; Newton et al., 2010; Spadone et al., 2015), but concentrates on tasks that share computational demands and overlapping topologies of evoked responses. In a sample of neurologically healthy adults (N=242), we first computed instantaneous functional correlation graphs, using a novel approach that temporally unwraps Pearson correlations to generate time series along edges, representing the inter-node BOLD signal co-fluctuations (Zamani Esfahlani et al., 2020). Then, by means of a general linear model (GLM), we assessed the task-based contributions to the edge time series, quantifying the amount of out-of-sample variability that they contained. We then compared the degree of between-task similarity at the regional activation and connectomic levels.

## Materials and Methods

### Participants

We analyzed task and resting-state fMRI data from the Pittsburgh Imaging Project (PIP), which is a registry of behavioral, biological and neural correlates of cardiovascular disease risk among otherwise healthy community-dwelling adults (aged 30–54 years). Details of this project can be found in the supplementary material of Gianaros et al. (2020). We selected a subset of 242 subjects (female=119, mean age=40 ± 6 years, min age=30 years, max age=51 years) that had full temporal and spatial coverage and exhibited low average motion (mean framewise displacement, estimated using the method in Power et al., 2012, lower than 0.35 mm) across the three fMRI tasks used in our study.

### MRI Data Acquisition

MRI data were acquired on a 3 Tesla Trio TIM whole-body scanner (Siemens, Erlangen, Germany), equipped with a 12-channel head coil. Functional blood-oxygen-level–dependent (BOLD) images were acquired from a T2*-weighted gradient echo-planar imaging sequence (repetition time=2000 ms, echo time=28 ms; field of view=205 × 205 mm (matrix size=64 × 64), slice thickness=3 mm (no gap); and flip angle=90°). For anatomical coregistration of the fMRI images, a high-resolution T1-weighted image per subject was also acquired (MPRAGE, repetition time = 2100 ms, echo time=3.29 ms, inversion time=1100 ms, flip angle=8°, field of view=256 mm × 208 mm (matrix size: 256 × 208), slice thickness=1 mm with no gap).

### Tasks

We used two tasks that involved processing conflicting information and response inhibition. Both tasks consisted of 4 blocks that defined a congruent information condition, interleaved with 4 blocks of trials where the participant received incongruent information. Both task conditions had a duration of 52-60 secs, and were preceded by a variable 10-17 sec fixation block. It total, each task had a duration of 9 min and 20 secs.

In the color-word Stroop task, participants had to select 1 of 4 identifier words using a response glove (e.g., thumb button 1 = identifier word on the far left, etc.), such that its name indicates the color of target words located in the center of a screen. During the congruent trials, the four identifier words were all in the same color as the target words. Instead, in incongruent trials, identifier words had all different colors, and the option to select was in a color incongruent with the target words. This kind of task usually evokes a brain response that activates regions in the anterior insula, parietal cortex, basal ganglia, thalamus, and cerebellum; while deactivating areas belong to the so-called ‘default-mode network’ (see Fig. 1A, B and C).

**Figure 1:**
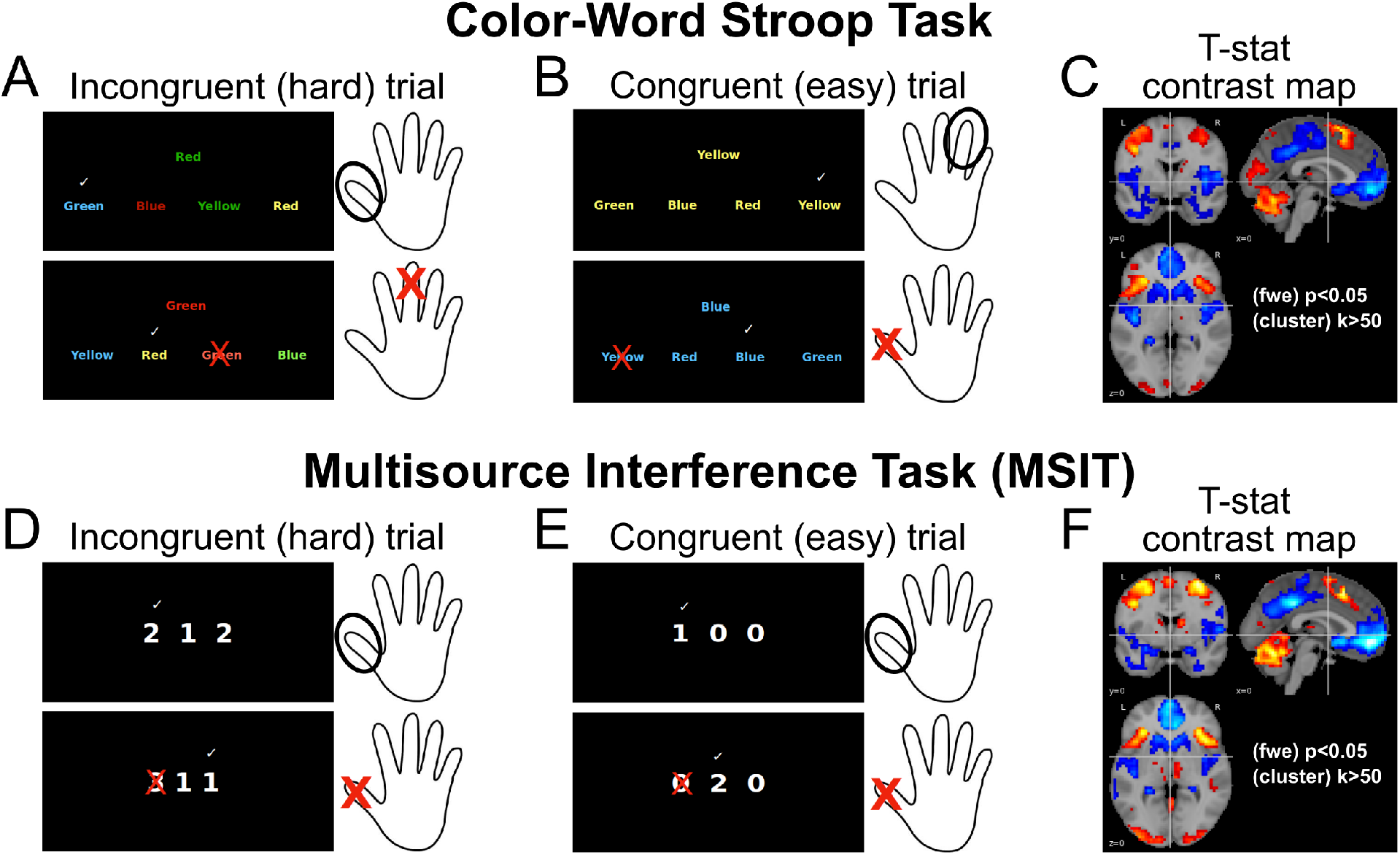
Stroop task and MSIT paradigms and their brain response. For both Stroop task and MSIT, illustration of incongruent (A, D) and congruent trials (B, E). Trials consisted of blocks of 52-60 s duration, interleaved with a 10-17 s fixation block. Contrasting brain activity between incongruent and congruent condition gives rise to a similar brain response (C, F).

In the MSIT, which corresponded here to a modification from the original task version (Bush and Shin, 2006), participants had to select 1 of 3 numbers such that it differed from the others 2 by pressing buttons on the glove, where each button matched a number on the screen (thumb button 1 = number 1, etc.). During congruent trials, targets’ position matched that on the glove, whereas during incongruent trials this position did not match the glove’s button location. This task elicits a brain pattern response that is largely similar to that in Stroop task (see Fig. 1D, E and F and Sheu et al., 2012 for more details on the MSIT and the Stroop task).

In incongruent conditions of *both* tasks, accuracy was titrated to ~ 60% by altering intertrial intervals, i.e. consecutive accurate choices led to shortened intertrial intervals. To control for motor response differences between conditions in both tasks, the number of trials in the congruent condition was yoked to the number completed in the incongruent condition. Yoking was implemented by (1) administering an incongruent block first and (2) presenting congruent condition trials at the mean intertrial interval of the preceding incongruent block.

Finally, we also used a five-minutes resting-state scan, during which the participants were told to keep their eyes open.

### Preprocessing

Data were preprocessed using fMRIprep (Esteban et al., 2018), a standard toolbox for fMRI data preprocessing that provides stability to variations in scan acquisition protocols, a minimal user manipulation, and easily interpretable, comprehensive output results reporting. First, anatomical data preprocessing was performed, including bias-field correction, skull-stripping, brain extraction and tissue segmentation, and surface reconstruction. It was then followed by functional data preprocessing, which included reference image estimation, head-motion parameters estimation, slice time correction, susceptibility distortion correction via a nonlinear registration (“Fieldmap-less” option of the toolbox), spatial normalization and confounds estimation.

### Functional Correlation (Edge) analysis

We estimated task-based functional correlations using the edge, or co-fluctuation, time series proposed in Faskowitz et al. (2020); Zamani Esfahlani et al. (2020). A sketch of this full estimation procedure can be found in Fig. 2. We first (step A) reduced the spatial resolution of the preprocessed time series by computing the voxel-wise average signal within each region (ROI) in a 268-parcels atlas (Shen et al., 2013). Following Finn et al. (2015), each of these regions were also identified to a specific intrinsic connectivity network: Motor, Visual-1, Visual-2, Visual-Association, Medial-Frontal, Frontoparietal, Default-mode and Subcortical-Cerebellum. Then, let 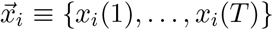 be the time series of *T* scans (the full-scan sequence) for a given parcel *i* in such atlas. Each of these parcellated time series were subsequently (step B) denoised by means of a linear regression model, in a single step that avoid artifacts from being reintroduced in the data (Lindquist et al., 2019), in order to remove effects from motion (24 parameters which included 3 translations, 3 rotations, their derivatives and the square of all these terms), the average white-matter signal, the average CSF signal, the average brain signal, periodic oscillations greater than 187 s (5 cosine terms) and task activations (24 terms). This last set of regressors consisted of 12 finite impulse response (FIR) terms per task condition (congruent and incongruent) to flexibly model an hemodynamic response function (HRF) of about 24s to external stimuli and that was included so as to avoid systematic inflation of functional correlations produced by task activations (Cole et al., 2019). The resulting denoised ROI time series were standardized (step C), i.e. 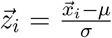, and then used to generate the edge time series 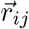 (step D) as the component-wise product between pairs of standardized time series, i.e. 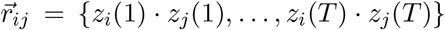. At this point, if we summed these components up and divide by *T* – 1, we would obtain the Pearson correlation coefficient that usually represents the static functional connectivity between BOLD time series - that is, each edge time series can be interpreted as a temporal decomposition of a functional connection (correlation) into its framewise contributions. Instead, we continued working on these edge time series as response variables in a general linear model (step E) in order to estimate intrinsic and task-dependent functional correlation profiles. To this end, the input design matrix included an intercept term and a set of regressors for each task condition (congruent and incongruent), which comprised a boxcar function convolved with the usual double gamma hemodynamic response function and its temporal and dispersion derivatives. Although it is not clear that the usual hemodynamic response function also takes place in the edge time series, we decided to assume it for pipeline compatibility with the activation (node) analysis (see next subsection). Yet, due to the nature of our tasks, we would not expect this to be a major issue, given that task-dependent GLM estimations essentially represented averages across blocks of considerable duration.

**Figure 2:**
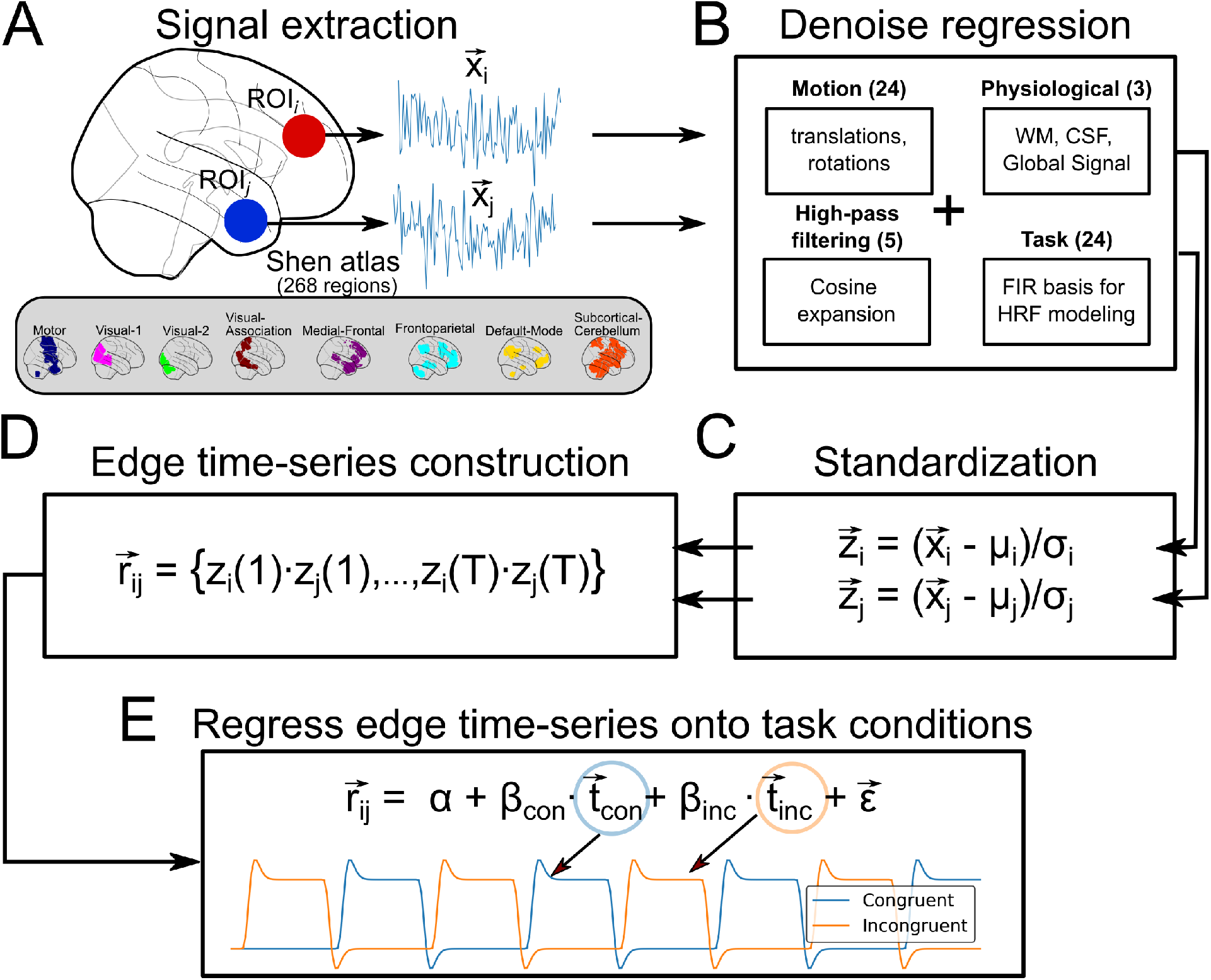
Estimation of intrinsic and task-related functional correlations. For a given pair of regions in the Shen atlas (consisting of 268 regions), the average signal within them was first computed (A). The time series were then denoised (B) and standardized (C). Subsequently, they were multiplied component-wise (D). Finally, the resulting temporal profile was regressed onto a design matrix to model intrinsic (intercept term) and task-related functional correlations (E).

Prior to any statistical analysis, time series in both sides of the regression model were prewhitened through a first-order autoregressive model in order to account for the temporal autocorrelations. As aforementioned, we assumed this standard procedure for dealing with temporal autocorrelations in order to have the same statistical pipeline as that in the activation analysis (see next subsection). Future studies should investigate the most appropriate procedure for accounting for autocorrelations using edge time series. After this first-level estimation, task-based network changes were computed as contrasts of parameters and subsequently used to assess edge-wise group-level effects by means of a one-sample t-test. Statistical inference at a usual 0.05 significance level was finally performed, after correcting the family-wise error (Holm-Bonferroni procedure) due to multiple testing. All these statistical analyses of the edge time series were carried out using Nilearn 0.7 (Abraham et al., 2014).

To test whether or not the inferred network maps reliably generalized to unseen data, the same general linear model was embedded within a k-fold cross-validation scenario. Specifically, for each edge time series used as a response variable and the task regressors (the same set of terms described previously) as predictors, the group-level *α* (intercept) and *β* (slopes) were estimated as their average across the first-level estimations from the subjects in the training set. Subsequently, these were used to quantify to what degree mean edge time series from the subjects in the test set were predicted. The difference between the predicted and observed edge time series was evaluated using the coefficient of determination *R*^2^. The advantage of using this metric is that for values greater than zero it means that the task stimuli explained a non-zero variability on top of that of the intrinsic or background signal, here modeled as the intercept. A 3-fold cross-validation was employed, where the full data were divided into 3 smaller sets (”folds”), such that for each of these (the test set) the generalization was determined using the coefficients estimated in the other 2 folds (the training set). This process was further repeated 20 times with different splitting seeds. The final performance per edge was then the average *R*^2^ across all folds and repetitions.

### Activation (Node) analysis

We also analyzed the preprocessed BOLD images at the node level, which involved estimating brain activation changes during the different task conditions. Such analyses are usually carried out at the voxel-level. However, in order to keep the same resolution as that of the edge-level results, brain activations were estimated at the region-level using the same parcellated BOLD time series.

For this analysis, we employed again a GLM with the parcellated BOLD time series as response variables and a design matrix that included the same set of task regressors used in the edge-wise analyses, as well as the same covariates that were regressed out prior to this, i.e. the 24 motion parameters (Friston et al., 1996), the cosine terms to account for oscillation effects greater than 187s, the average signal within whitematter tissue, the average signal within CSF tissue, and the average signal within the whole-brain. We considered this last regressor, not common in brain activation analyses, for consistency again with the edge-level analyses (see previous section). Group-level effects were similarly assessed using a one-sample t-test.

### Generalized Psychophysiological Interaction

As a part of our sanity check pipeline, we compared the functional correlation analysis using the aforementioned edge time series approach with a model of Generalized Psychophysiological Interactions (PPI), which is a standard approach for estimating task-dependent functional connectivity changes (McLaren et al., 2012) and is based on a general linear model of task-moderated temporal association between pairs of brain units. Specifically, for a given pair of BOLD time series 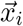 and 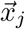, such model includes one of them as the response variable and as inputs the other BOLD time series, the group of task regressors, the interaction terms between these task regressors and the input BOLD time series, and the possible confounders to consider in the model. In our case, the generalized PPI model can be written as follows:

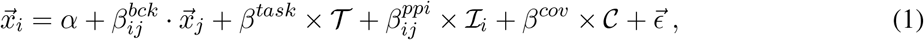

where 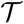 is a matrix whose columns are the HRF convolved box-car congruent and incongruent time profiles and their derivatives and dispersion terms, 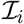 the matrix with the PPI terms from each condition, i.e. the interaction term between each task condition’s time profile and the input time series 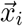, and 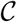 a matrix with the different covariates to include in this model, which in our case comprised the 24 motion parameters, the average white-matter signal, the average CSF signal, the average brain signal and cosine expansion for a 187 sec high-pass filtering. Once all the parameters in this model were estimated, task-based connectivity changes were evaluated by contrasting the incongruent and congruent PPI estimations and their effect at the group level assessed using a one-sample t-test. In this way, a matrix of estimated task-based connectivity changes can be constructed. However, since a PPI model yields non-symmetrical matrices, we symmetrized them by averaging their corresponding upper and lower triangular elements as done in Di et al. (2017), which enabled a direct comparison with the connectivity profiles obtained from the edge time series approach.

## Results

### Group-level activation patterns

We begin by replicating an exhaustively reported effect (see Sheu et al., 2012 and references in), namely that the Stroop task and MSIT, both effortful cognitive control tasks, have largely overlapping spatial patterns of evoked activity across the brain, particularly the neocortex (see contrasts maps in Fig. 3). Here such similarity was quantified by a Spearman correlation coefficient, *ρ*, between un-thresholded incongruent-vs-congruent t-stat maps calculated at the region-level (voxel-wise estimations with the same region-size spatial smoothing yielded similar values), and a Dice similarity coefficient (DSC), from binarizing these maps as to whether their t-stats rejected or not the null hypothesis at *α* = 0.05 after family-wise (Holm–Bonferroni) error correction. For our group-level activation patterns, the former, *ρ*, was equal to 0.87, and the latter, DSC, equal to 0.86. As shown in Fig. 3, increases in brain activity in incongruent trials, with respect to congruent trials, were located in areas typically engaged during the processing of conflict information and response inhibition, such as the anterior cingulate cortex, anterior insula, parietal cortex, basal ganglia, thalamus, and cerebellum. In contrast, de-activations took place in areas within the ventromedial prefrontal cortex, perigenual anterior cingulate cortex, posterior cingulate cortex, and precuneus, which all comprise the default-mode network. As a consequence, these results show that similar cognitive contexts evoke similar patterns of activity outputs in areas segregated across the brain.

**Figure 3:**
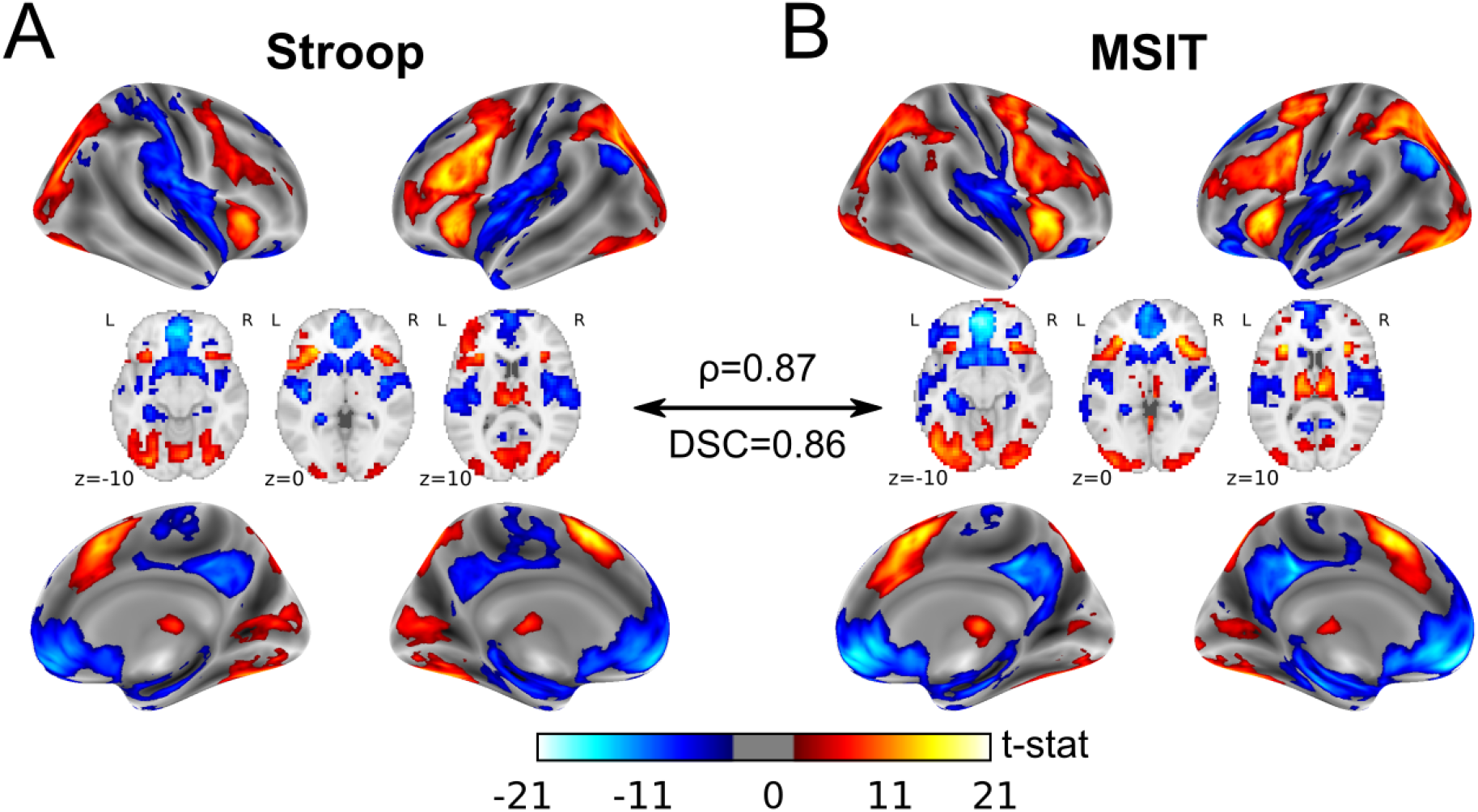
Group-level activation maps. For both Stroop task (A) and MSIT (B), the group-level incongruent-vs-congruent t-stat maps, at the voxel level for aesthetic reasons. Thus, red colors display higher BOLD activity during incongruent trials compared to congruent trials, whereas blue colors represent the other way around.

### Exploration of co-fluctuating hemodynamics

For illustrative purposes, we examined the task-related effects on the inter-region co-fluctuations by computing the root sum of squares (RSS) across edges at each time frame. It is important to clarify that, for this calculation, parcellated BOLD time series prior to edge time series formation included all task events, in contrast to subsequent analyses. As shown in Fig. 4A and B, during both tasks high moments of co-fluctuations tended to be synchronized across subjects, concentrated mostly around the rest periods separating congruent and incongruent block conditions. In both the congruent and incongruent blocks, there appeared to be a consistent reduction in global connectivity, with sporadic and inconsistent periods of brief synchronous activity that qualitatively appear more frequent during incongruent blocks.

**Figure 4:**
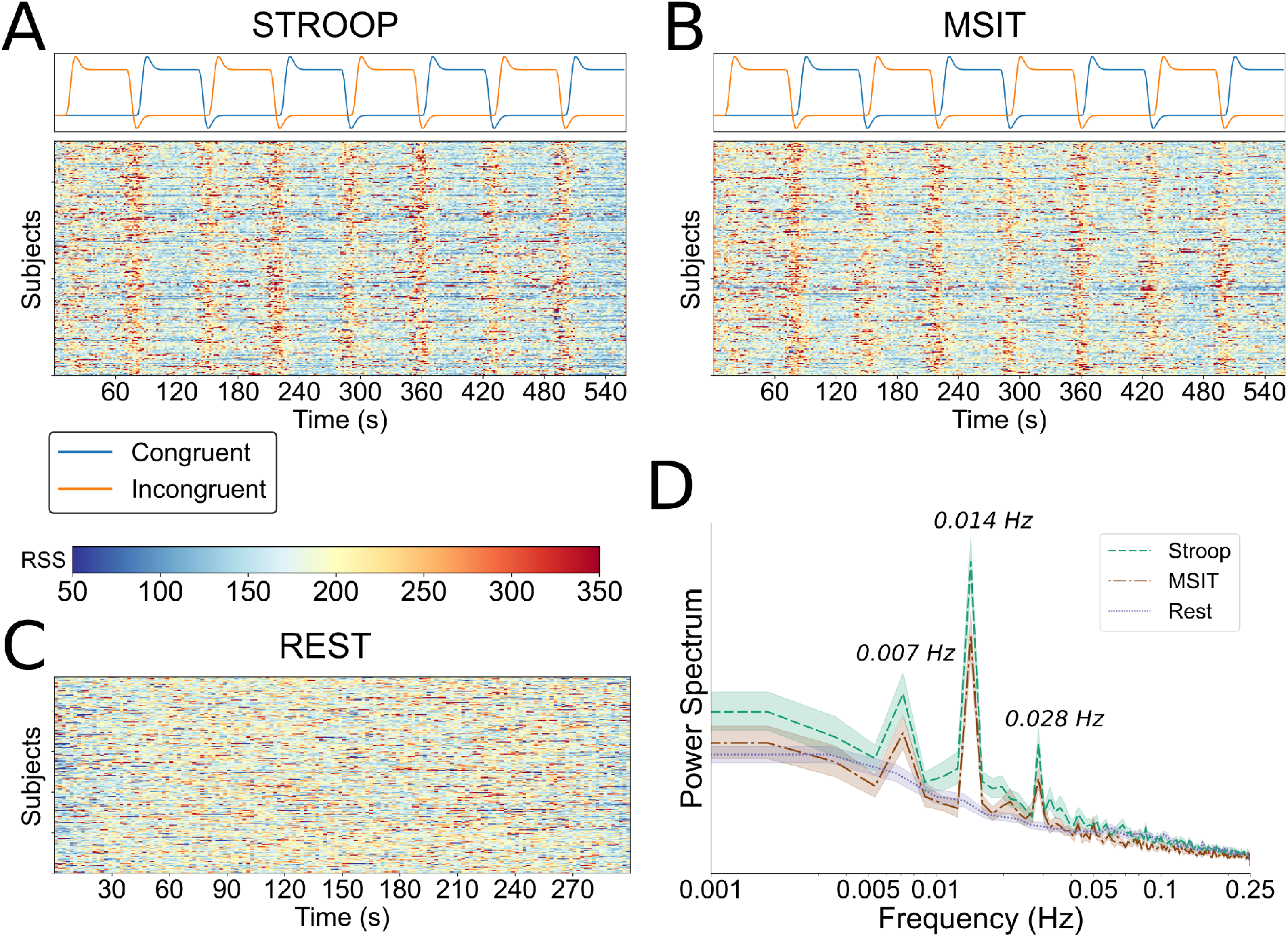
Analysis of the root sum of squares time series. For each subject, the root sum of squares of the edge time series that include the task effects for Stroop task (A), MSIT (B) and resting-state acquisition (C). Their power spectrum (in arbitrary units) using a periodogram (D), averaged across subjects.

In contrast to the task patterns, for the resting-state run, where no external stimulus was presented, we did not see evidence of between-subjects synchronization of high amplitude co-fluctuations (Fig. 4C). Though the overall presence of these brief co-fluctuations appears to be more frequent in the resting-state run than during either of the two tasks. These results were further confirmed by inspecting the subject-averaged power-spectrum of the RSS for the three tasks (Fig. 4D). For both Stroop and MSIT, there was an overall increase in power at frequencies consistent with task onsets and offsets.

### Group-level functional correlations

We estimated task-related functional correlations using a GLM on the edge time series. Three coefficients (i.e., intercept, congruent, and incongruent) for both Stroop task and MSIT were estimated for each edge, while for resting state a single coefficient per edge was obtained (i.e., intercept only). The resulting group-level network profiles are displayed in Fig. 5, where the t-stats for each of these coefficients were converted to correlations using the transformation 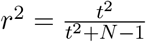, with N being the number of subjects.

**Figure 5:**
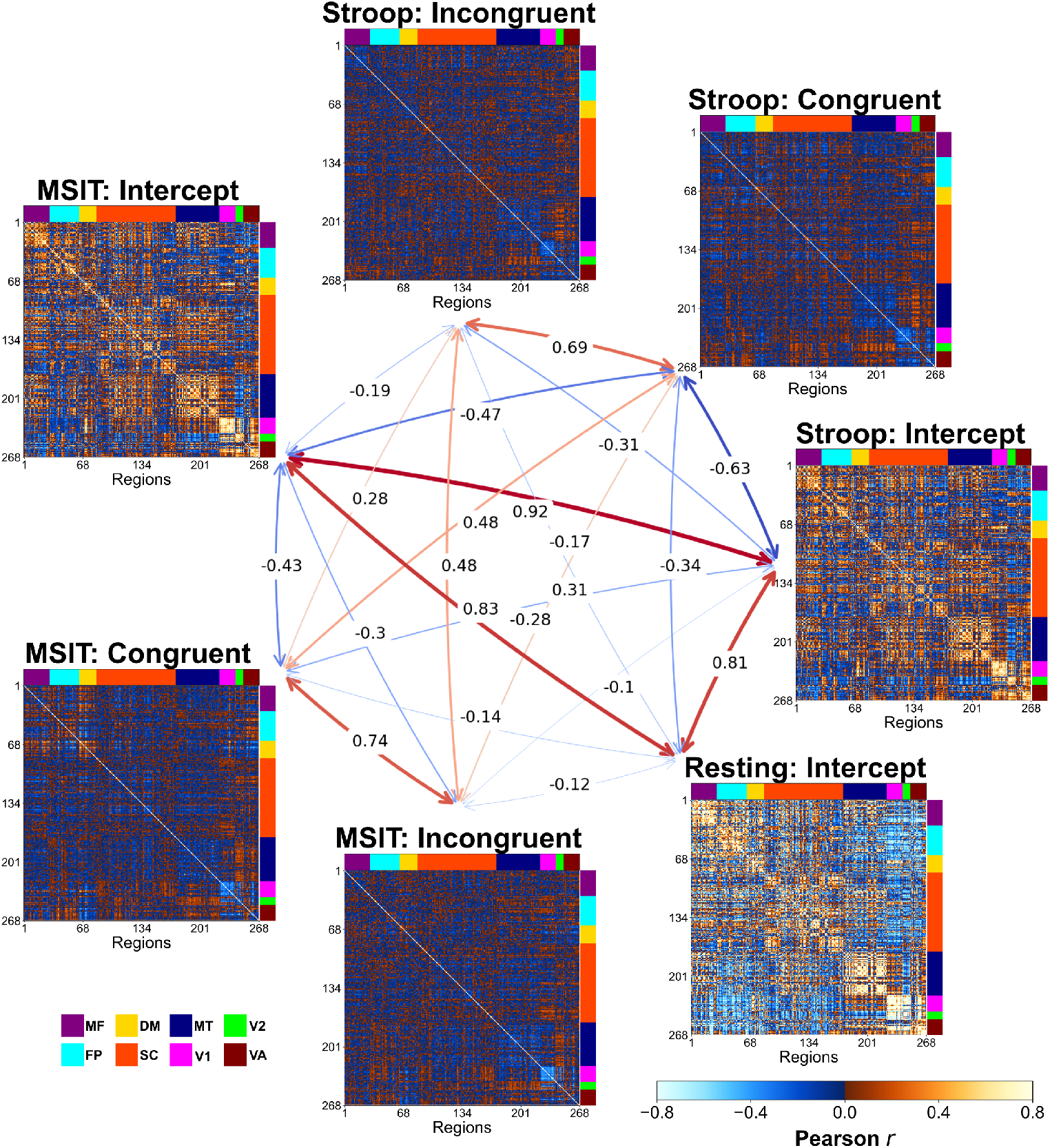
Functional correlation matrices at the group level. For Stroop task, MSIT and resting-state functional correlation matrices using the intercept, congruent and incongruent GLM estimations at the group level. Regions (i.e. the rows and columns) have been arranged based on their belonging to a major intrinsic network system (see Methods). In the middle in the form of a graph, the Spearman correlations between the upper-triangular elements of these matrices. MF: Medial-Frontal; FP: Frontoparietal; DM: Default-mode; SC: Subcortical-Cerebellum; MT: Motor; V1: Visual-1; V2: Visual-2; VA: Visual-Association.

The first thing to note is that, after accounting for condition effects during the two tasks, we were able to recover the intrinsic brain networks observed during resting state. The intercept profiles for both Stroop and MSIT had a high degree of similarity to the resting state profile (*ρ* = 0.81 and *ρ* = 0.83 respectively), as well as a high degree of similarity to each other (*ρ* = 0.92).

On the other hand, a largely different profile emerged during congruent and incongruent conditions in both tasks. These networks showed much lower overall functional correlations, and a shift towards more negative correlations, than the intercept profiles. Despite this difference from the intrinsic networks, the condition-related profiles (i.e. congruent and incongruent) had a decent degree of within-task similarity (*ρ* = 0.69 for Stroop and *ρ* = 0.74 for MSIT), demonstrating that both conditions recruit largely consistent networks overall. Less similarity was observed between-task profiles, whether it be using within condition comparisons (*ρ* = 0.48 in both cases), or between-condition comparisons (Stroop Congruent-MSIT incongruent ρ = 0.31, Stroop Incongruent and MSIT Congruent *ρ* = 0.28).

Taken together, these results confirm that our method was able to reliably characterize both task and intrinsic (resting) networks, at the group level, using the edge time series.

### Network profile differences between task conditions

The network profiles that emerged as a consequence of conflict processing were quantified at the group level by contrasting subject-level functional correlations from both task conditions. The resulting incongruent-vs-congruent statistical maps for both tasks are displayed in Fig. 6 (left plots, panels A and B), with 1228 (Stroop task) and 1076 (MSIT) edges that were significant at *α* = 0.05 after family-wise (Holm-Bonferroni procedure) error correction (red colors denote a greater functional correlations during incongruent trials than during congruent trials, and blue colors the opposite). In both cases, network differences were primarily associated with default-mode, frontoparietal, medial-frontal and visual systems, as measured by the average significant edges per region found in those networks. Furthermore, inspecting the sign of these differences (Fig. 6, right side of panels A and B), increased functional correlations appeared to be dominated by edges connecting regions of distinct intrinsic major systems, particularly those between the default-mode and the frontoparietal and visual-association systems, and medial-frontal areas with the frontoparietal cortex. In contrast, significant decreases in functional correlations during incongruent trials appeared in regions of the same major system, specially those within the default-mode and medial frontal networks.

**Figure 6:**
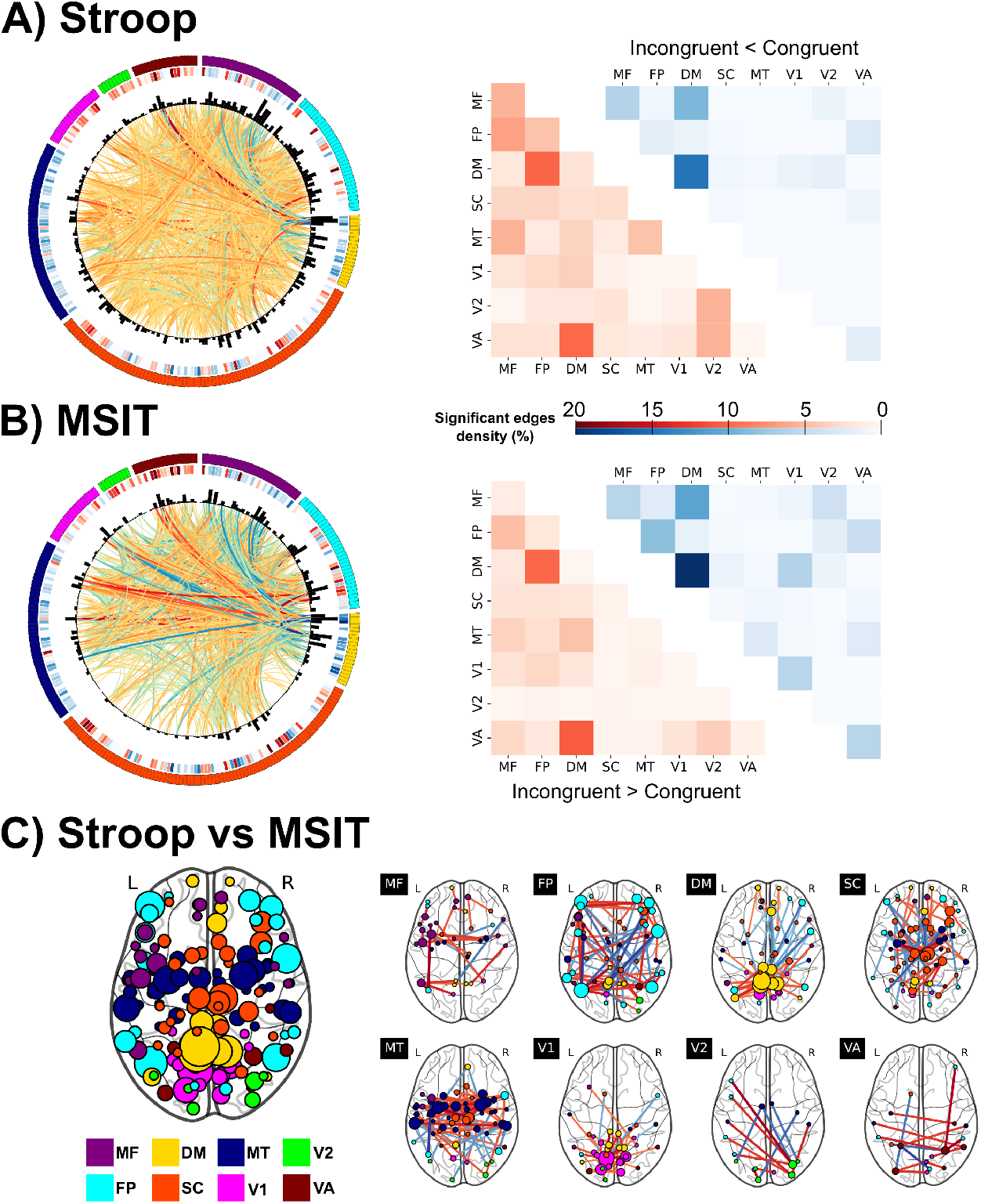
Group-level incongruent-vs-congruent functional correlation differences. For Stroop task (A) and MSIT (B), on the left side and from outer to inner circular, plots display each region arranged and colored according to the major functional system, their incongruent-vs-congruent activity at the node level, their degree from the incongruent-vs-congruent significant edges, and finally the t-stat of these edges (red: incongruent > congruent, blue: incongruent < congruent). At both node and edge-level, only significant results (at *α* = 0.05, Bonferroni corrected) are shown. On the right side the number of significant edges within and between major connectivity networks, normalized by the total number of edges in each case. (C) Using the significant edges from a paired t-test at *α* = 0.05 Bonferroni corrected, between-task differences in incongruent-vs-congruent functional correlations shown in region degree (left panel), and with the edge t-stats to regions of each major functional system (right panel). MF: Medial-Frontal; FP: Frontoparietal; DM: Default-mode; SC: Subcortical-Cerebellum; MT: Motor; V1: Visual-1; V2: Visual-2; VA: Visual-Association.

However, despite the apparent qualitative similarity in network-level responses to congruent and incongruent conditions, the Stroop and MSIT also exhibited key differences. For example, concentrating on the 10% of edges with the largest absolute t-stat values (n=358), the Stroop task contained a significantly greater number of positive (i.e. increased functional correlation during incongruent trials) to negative (i.e. decreased functional correlation during incongruent trials) edges than the MSIT (Fisher exact’s test, odds ratio = 3.76, *p* = 4.00 × 10^-14^). On the other hand, a paired-sample t-test performed on individual edges revealed that these between-task network differences spanned the entire brain (see Fig. 6C), though they prominently expressed in the dorsolateral prefrontal and posterior parietal cortex, both responsible for executive function, as well as in the posterior cingulate cortex, that is strongly implicated during control processes, and the primary visual cortex. As a consequence, these results suggest that the Stroop task and MSIT have substantial differences in their network profiles.

### Reliability of edge-wise responses

We next explored the reliability of functional correlation graphs shown by testing whether edges could be predicted out-of-sample by the tasks. Since we had two fMRI tasks, this gave rise to two possible predictive scenarios: “within-tasks”, if out-of-sample predictions were performed on the same task used for training, or “between-task”, if out-of-sample performance was tested on the task not used for training. The edge-wise prediction rates are displayed in Fig. 7A and B, contrasting within- and between-task performance for both Stroop and MSIT data used as the training set. First, it is noteworthy that approximately 88% (Stroop) and 94% (MSIT) of the significant edges resulting from the incongruent-vs-congruent contrast reported in previous sections could be predicted here within-tasks, demonstrating the reliability of these network profiles. Second, one can see that the task-based signal at each edge varied dramatically across the entire brain network, with some edges containing more than 30% variability attributable to the task conditions. Nevertheless, across both Stroop and MSIT only less than 10% of the edges appeared to encode meaningful task variability (i.e. average *R*^2^ greater than zero at 95% confidence level).

**Figure 7:**
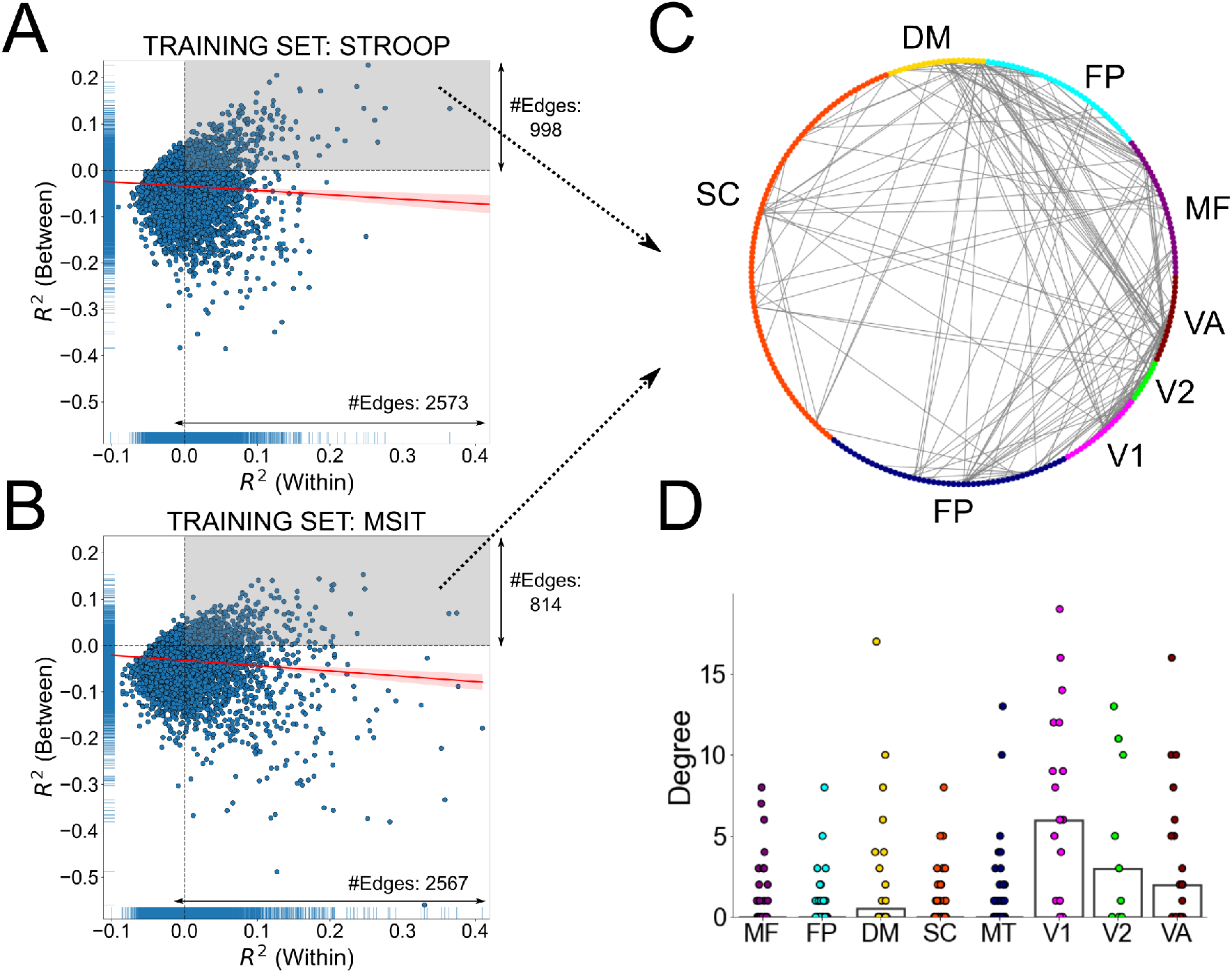
Edge-wise out-of-sample generalizability. The coefficient of determination for each edge predicted by the tasks regressors, using A) Stroop and B) MSIT as training set. Values on the horizontal axis represent predictions on the same task used for training (within-tasks), whereas between-task values (predictions on the different task used as training) lie on the vertical axis. C) Binary graph that displays the sub-network formed by those edges that could be predicted (average *R*^2^ > 0 at 95% confidence level) within and between-task in both MSIT and Stroop task (points in the gray area in panels A and B). D) Using the aforementioned sub-network, degree (i.e., number of edges connected to each node), grouped by major macroscopic functional systems. Each point then represents a region. The bars show the median degree across regions within each intrinsic network. MF: Medial-Frontal; FP: Frontoparietal; DM: Default-mode; SC: Subcortical-Cerebellum; MT: Motor; V1: Visual-1; V2: Visual-2; VA: Visual-Association.

More importantly, edges largely tended to generalize better within-tasks. For example, during the Stroop task 2573 edges could be predicted if tested on the same task (median *R*^2^ = 0.026 across these edges), whereas between-task this number dropped to 998 (median *R*^2^ = 0.022 across these edges). Moreover, only 474 of these edges could be predicted successfully both within and between-task, so the intersection between binary masks of predicted edges in both cases was low (*DSC* = 0.265). Likewise, similar values were found for the MSIT (2567 predicted edges within-tasks and median *R*^2^ = 0.028 across them, 914 predicted edges between-task and median *R*^2^ = 0.016 across these edges, intersection *DSC* = 0.228). Consequently, within- and between-task predictions had a poor correspondence (see the almost flat regression line in Fig. 7A and B), which is consistent with the differences in network profiles between both the MSIT and Stroop task observed in previous sections. Interestingly, in both scenarios there was a subset of edges that were not predicted by the same task used for training, but by the other task (points in the left upper corner of Fig. 7A and B). However, the *R*^2^ values for these are quite small and likely reflect random noise in the generalization error estimates.

Finally, we wondered which sub-network(s) appeared to be most consistent across tasks by looking at the shared edges between the Stroop and MSIT that could be predicted both within *and* between-task (points between the right upper corners of Fig. 7A and B). The resulting sub-network contained 239 edges (Fig. 7C), with a prominence of connections to areas within and between visual systems (see Fig. 7D), suggesting that part of the visual information is similarly encoded and preprocessed by both tasks, and some areas in the default-mode network.

### Comparison of similarities in activation patterns and network profiles between tasks

We have previously shown that both Stroop and MSIT elicit largely overlapping patterns of brain activation (*ρ* = 0.87, *DSC* = 0.86; see also Sheu et al., 2012). In contrast, estimated edge-wise responses suggest that both tasks appeared to differ at the network level. Is the lower similarity of network profiles between-task really that different than the similarity in activation patterns? The between-task similarity in incongruent-vs-congruent network profiles was equal to *ρ* = 0.64 and *DSC* = 0.43 at *α* = 0.05, after family-wise (Holm-Bonferroni) correction, which indeed constitute a considerable reduction with respect to the aforementioned similarity rates from activation patterns. Furthermore, this reduction became even more evident as the number of subjects decreased (Fig. 8A), suggesting that this does not reflect an issue with statistical power in our sample. Also, this effect is largely insensitive to using Spearmans *ρ* as a similarity measure since the same effect was observed using Dice similarity coefficients at different thresholds (see Fig. 8 A and B).

**Figure 8:**
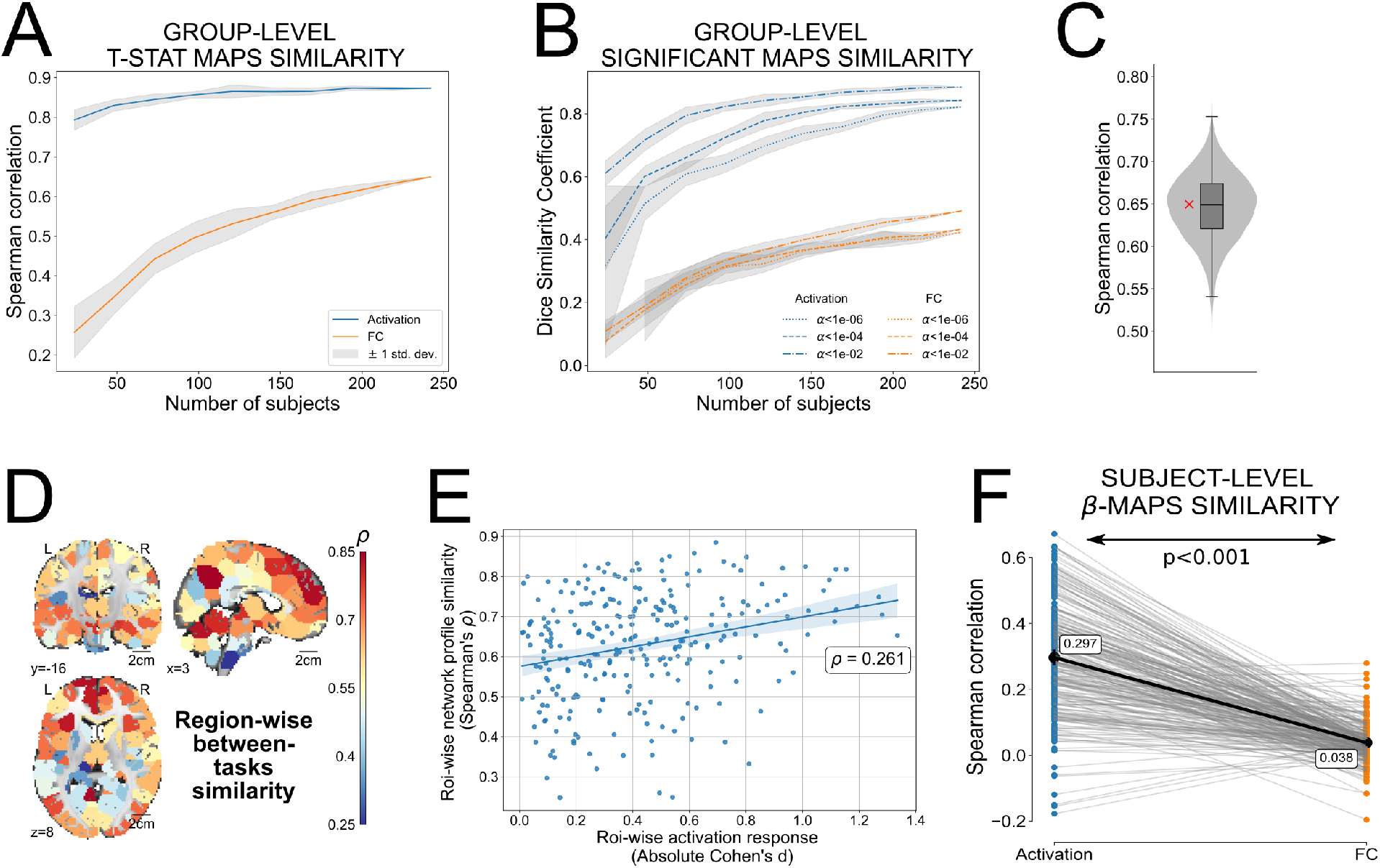
Between-task similarity of activation patterns and network profiles. A) Spearman correlations between tasks from the group-level t-stat incongruent-vs-congruent maps for both brain activation (blue line) and task-based functional correlations (FC, orange line), varying the number of subjects used for their estimation. Each curve represents the average similarity and the gray area the standard deviation after repeating 10 times the estimation procedure to consider different subjects. B) Same as A) but using the dice similarity coefficient. Statistical maps were binarized according to whether each t-stat was significant or not under several thresholds *α*. C) Distribution of Spearman correlations *ρ* between MSIT and Stroop functional correlation profiles from 10000 subsamples that each randomly selected a subset of edges equal to the number of regions (268). The red cross displays the correlation using the full profiles (i.e. 35778 edges). D) Region-wise similarity between tasks, using the whole-brain incongruent-vs-congruent network profile of each region. E) These similarity rates per region (y-axis) are plotted versus their activation levels, measured as the average of both tasks’ incongruent-vs-congruent absolute Cohen’s d at the group level. F) For each subject (a dot in the figure), the Spearman correlation between the incongruent-vs-congruent *β* map of each task for both brain activation (blue points) and task-based functional correlations (orange points). A paired t-test then quantified the statistical difference between both distributions.

In order to show that this reduction in similarity scores between the incongruent-vs-congruent functional correlation graphs was not due to correlating a larger number of features from the edges (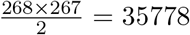 edges) than in the activation maps (only 268 components, since these were also considered at the region-level), we repeated this calculation taking subsamples (number of subsamples=10000) that randomly selected 268 edges in the functional correlation profiles. Across all subsets, we found similar between-task Spearman correlation values (0.65 ± 0.04, see Fig. 8C) as the one using the full network.

Along similar lines, we explored how similarity of network profiles was expressed across the brain by correlating, for each region, the whole-brain incongurent-vs-congruent functional correlation profile (a vector of 267 t-stat values, i.e, we do not consider the diagonal terms in the functional correlation profiles) at the group level of both tasks (see Fig. 8D). This analysis showed that there are certain regions, particularly in the superior medial and dorsolateral-frontal gyrus, the precuneus and the anterior lobe of cerebellum, that exhibit comparable, and sometimes even greater, similarity values than that from activation patterns. While regions with the largest between-task similarities in activation did tend to have higher degrees of between-task similarity in network profiles (Fig. 8E), this association was fairly weak (*ρ* = 0.261), suggesting that our main conclusion would also be reached if one focused exclusively on the sub-network typically engaged during both Stroop and MSIT.

Since the previous calculation concentrated exclusively on group-level patterns, we also tested whether the same qualitative findings were present at the within-subject level. Specifically, for each individual we correlated, between tasks, the incongruent-vs-congruent activation maps and functional correlation graphs, using in both cases the *β* estimations (see the sample distributions in Fig. 8F). The reason for using the *β* estimations here instead of the t-stat values is that temporal autocorrelations in the time series produced a different number of degrees of freedom across nodes and edges in both tasks, in contrast to the group level, where the degrees of freedom remained always the same (*N* – 1, with *N* the number of subjects). A paired t-test showed that, as found before with the group-level maps, between-task similarity rates of brain activation maps (mean *ρ* = 0.30, 95% CI [0.279, 0.322]) were higher than those from task-based network differences (mean *ρ* = 0.038, 95% CI [0.031 0.046]; Cohen’s *d* = 2.07, *p* < 0.001).

We ran several follow up tests to ensure that all these findings did not depend on choices taken during the setup of the analytical pipeline. First, we investigated whether the reduction in between-task similarity values from network profiles compared with evoked responses was not due to regressing out the task stimuli prior to the calculation of the edge time series. As shown in Fig. 9A, a mild increase was obtained when tasks effects were maintained (mean *ρ* = 0.075, 95% CI [0.066, 0.083]) and it was still significantly lower with respect to activation patterns (Cohen’s *d* = –1.71, *p* < 0.001). Moreover, a similar finding was observed (see Fig. 9B) when we concentrated exclusively on the regions with greatest task activation response (group-level incongruent-vs-congruent absolute Cohen’s d’s larger than 0.8 in both tasks), given that these regions should be particularly sensitivity to the removal of task stimuli. Thus the choice of regressing out task effects prior to building the edges-time series did not drive our primary effect.

**Figure 9:**
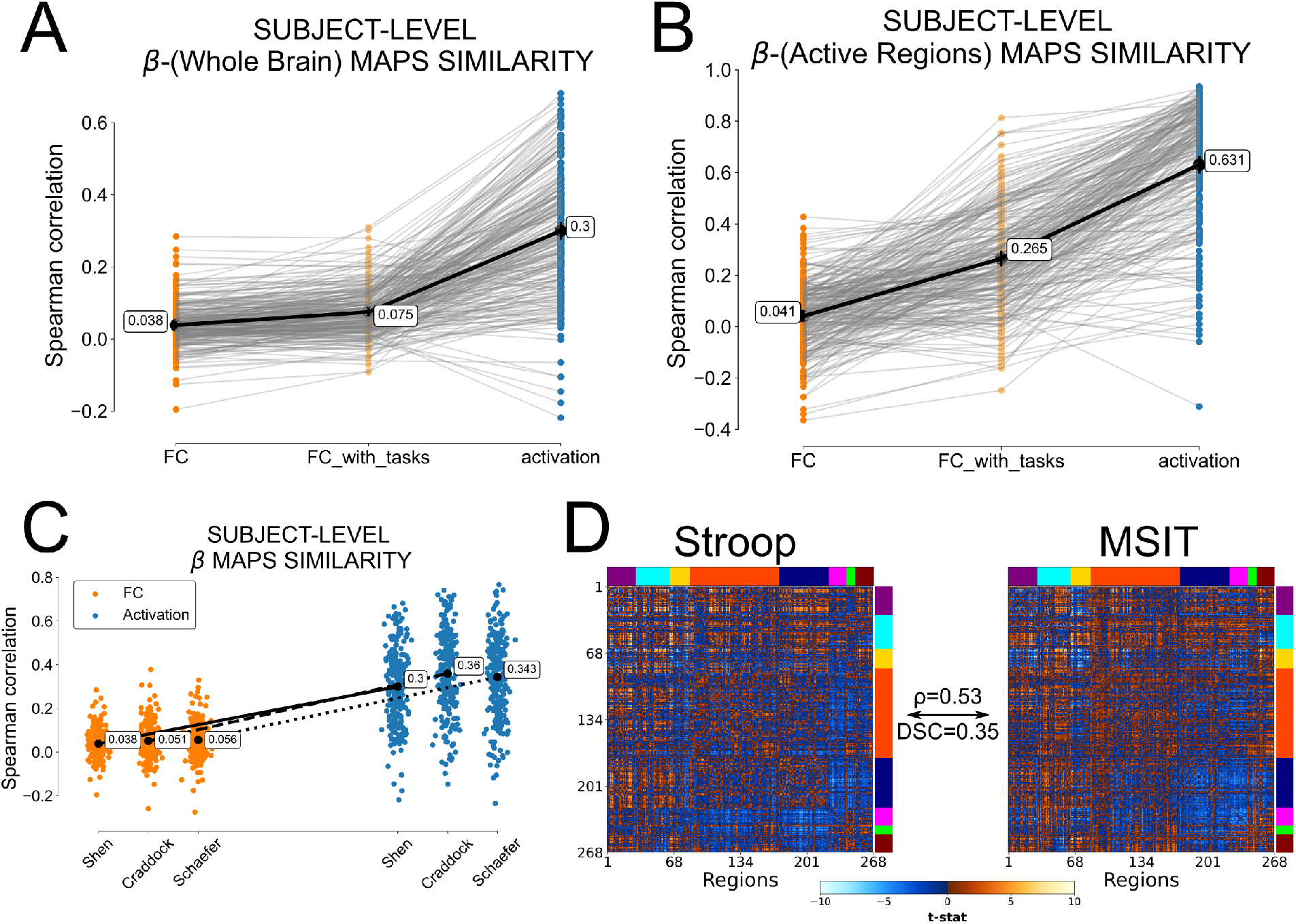
Replication analyses. A) Same subject-level results as in Fig. 8F, but including the case where task stimuli were not regressed out prior the edge time series formation (”FC with tasks”). B) Same as A), but concentrating on the subset of regions with greatest task evoked response (group-level incongruent-vs-congruent absolute effect sizes, i.e. Cohen’s d, in activation larger than 0.8 in both tasks) C) At the subject level, between-task Spearman’s *ρ* correlation from brain activation (blue points) and task-based functional correlation (FC, orange points) *β* profiles for Shen, Craddock and Schaefer parcellations. D) PPI-based incongruent-vs-congruent connectivity t-stat matrices for both Stroop and MSIT, and the similarity (Spearman’s *ρ* and DSC) between them.

Subsequently, we tested whether including in our parcellation specific brain structures that are known to be noisier or more susceptible to signal loss, namely the cerebellum and subcortex, might have driven our findings. In order to achieve this, we repeated the subject-level similarity analysis using a Craddock atlas (Craddock et al., 2011), consisting of 200 regions that did not include the cerebellum, and the Schaefer atlas (Schaefer et al., 2017), comprising 200 cortical regions. We found, again, that the choice to use the Shen atlas was not decisive in our primary effect of the differences between activation and network profile similarities (See Fig. 9C).

Likewise, we wondered whether the reduction in similarity between Stroop and MSIT task-dependent network profiles was influenced by the edge time series approach itself. We tested this possibility by replicating our analyses using a generalized Psychophysiological Interaction (PPI) model, which is a standard and common framework for assessing task modulated functional connectivity (see Materials & Methods for details on this model). As Fig. 9D illustrates, these PPI-based network profiles also showed a reduced similarity between tasks (*ρ* = 0.53, *DSC* = 0.35) compared to what is observed in the brain activation patterns. In addition, albeit small differences existed, particularly within the motor system, both approaches (edge time series and PPI) appeared to yield fairly similar incongruent-vs-congruent contrast network profiles in both tasks (*ρ* = 0.74 for Stroop, *ρ* = 0.77 for MSIT).

Finally, due to the ongoing controversy around the inclusion of the brain global signal in task-effect estimations (Liu et al., 2017), we recomputed both incongruent-vs-congruent activation maps and functional correlation graphs without the global signal removed. We found even more pronounced topological differences in network profiles between Stroop and MSIT with the global signal left in, which translated in the same reduction in group- and subject-level between-task similarity rates compared to those from activation patterns (see Fig. 10). Therefore, our difference between activation and network profile similarities cannot be explained by the absence of the global signal.

**Figure 10:**
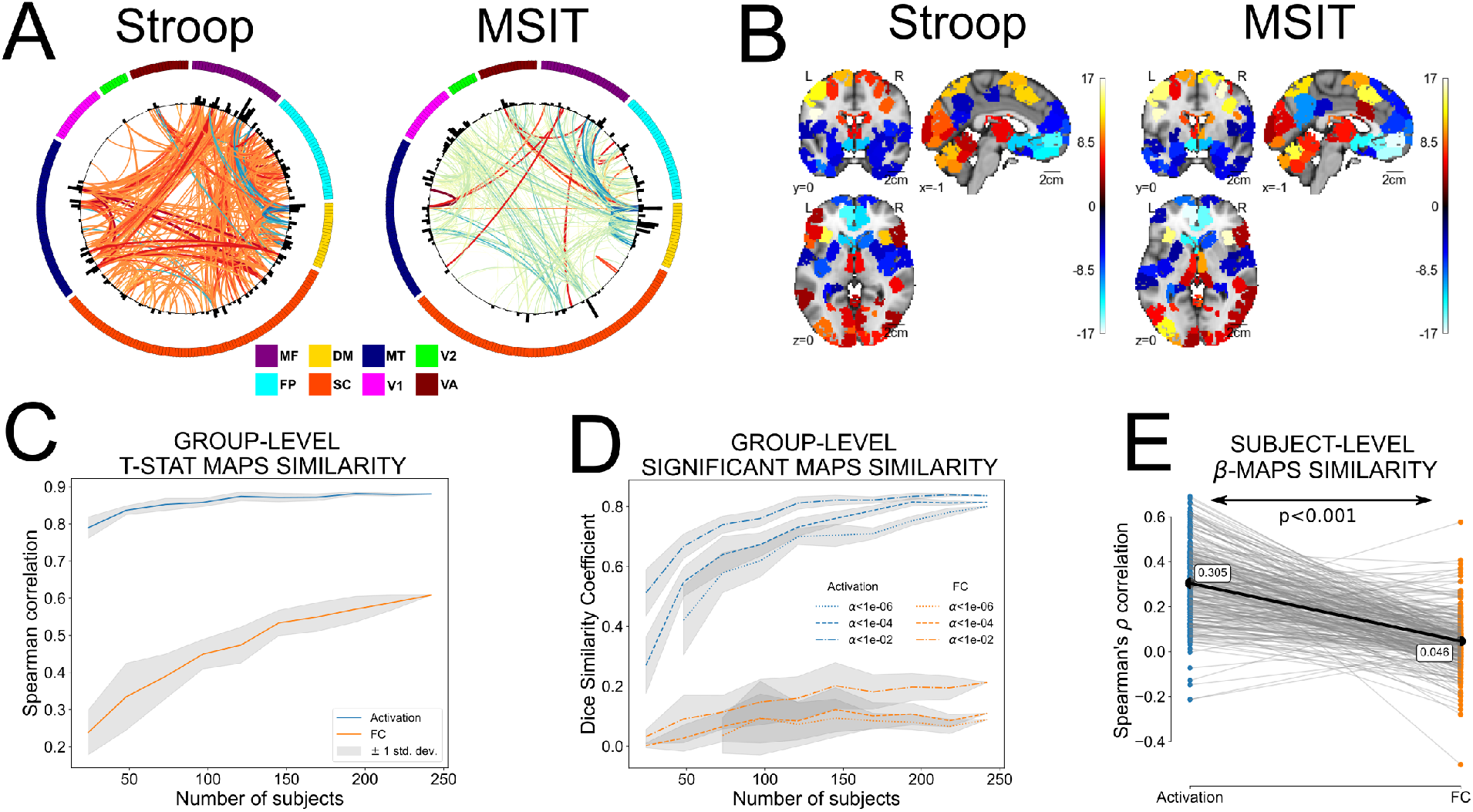
Results with global signal. A) Network profiles as incongruent-vs-congruent differences for both tasks. Each edge depicts the t-stat value of this contrast at the group-level (red: incongruent > congruent, blue: incongruent < congruent) B) Same as A) but depicting instead brain activation patterns as incongruent-vs-congruent t-stat contrast maps E) Spearman’s *ρ* correlations between tasks for both activation (blue line) and functional correlation profiles (FC, orange line) at the group-level, varying the number of subjects used for their estimation. Each curve represents the average similarity and the gray area the standard deviation after repeating 10 times the estimation procedure to consider different subjects. F) Same as E) but using instead a dice similarity coefficient between binarized significant maps for different thresholds. G) Spearman’s *ρ* correlation between subject-level incongruent-vs-congruent *β* profiles of each task for both brain activation (blue points) and task-based functional correlations (orange points).

## Discussion

Here we set out to determine whether two tasks with highly similar activation patterns also share common task-related network profiles. Using a GLM framework on instantaneous functional correlation estimates (Faskowitz et al., 2020; Zamani Esfahlani et al., 2020), we were able to successfully separate task-free (intrinsic) from task-dependent network contributions, in line with the extensive evidence that task functional correlations are jointly shaped by both intrinsic and evoked network architectures (Cole et al., 2014). Subsequently, we showed how our two tasks shared a large degree of similarity in activation topology (nodes), but substantially less similarity in network profiles (edges). This difference in task effects at the nodes and edges was confirmed at both group and subject level and using two different measures commonly employed for representational similarity analyses. Likewise, this difference between activation and network profiles was replicated after keeping task effects in the edge time series, employing different parcellations, using a different method for estimating task-related connectivity (i.e., PPI), and without including the brain global signal as covariate. Taken all together, these results are consistent with the dynamical systems perspectives of the macroscopic brain that suggest tasks are separately represented at both the node (voxel or region) and edge (connectivity) levels.

How is it possible that similar patterns of brain activation arise from dissimilar network architectures? Even though our study did not enable us to directly address this question, our results provide intriguing hints that the underlying mechanisms responsible for this take place at the connectivity level. It is important to keep in mind that functional connectivity is, itself, a dynamic signal. Even though intrinsic networks, like those measured in resting-state conditions, are thought to reflect underlying structural (or at least static) networks (Hagmann et al., 2008; Honey et al., 2009; Cole et al., 2014, 2016), the reality is that correlations in the hemodynamic BOLD response reflect highly flexible network states. Indeed, it is well known that these networks can flexibly reconfigure in order to adapt to contextual changes (Spielberg et al., 2015), so one could expect a similar behavior even when these contextual changes do not produce a significant deviation in the brain response. In our case, network differences between tasks are not large (i.e., they are not completely independent networks), albeit significantly larger than those from brain activation patterns. Nevertheless, it is expected that these differences become more pronounced as the scale of the interacting components increase, e.g., at the neuronal level (Prinz et al., 2004; Hooper, 2004; Cropper et al., 2016).

It is worth pointing out that while network profiles do differ more between tasks than activation patterns, we still observed a modest degree of similarity in network profiles across tasks. This is not surprising given the existence of a core functional architecture shared between even markedly different task states (Krienen et al., 2014). In our case, the greatest similarities were found in networks that are reliably associated with sensory processing and motor planning. While motor planning constraints were identical across tasks (i.e., both involved button presses with the same hand and fingers), the visual stimuli were quite different (see Fig. 1). This suggests that the between-task dissimilarities in network profiles reflect differences in *how* sensory information is used during action selection, after sensory representations are formed, rather than simple bottom-up effects driven by the stimulus differences between the Stroop task and MSIT. Adding to the other between-task topological differences that we observed, involving mainly regions of the defaultmode and executive networks, this appears to suggest that greater deviations take place in subnetworks largely associated with higher-level cognitive functions.

One natural follow-up question is how the edge time series responses compare to other approaches for addressing task-based networks like PPI. We have shown that, even though PPI arrives at the same conclusion as the edge time series method, the network profiles obtained from both approaches were not perfectly identical. While the edge time series straightforwardly represents measures of (instantaneous) functional correlations, PPI was designed to assess effective connectivity (Friston et al., 1997; Friston, 2011). Thus, in order to enable the comparison between both approaches in our study, PPI estimates were symmetrized, so we speculate that part of these differences may come from this operation. A full comparison with other common methods for task-related networks, such as correlational PPI (Fornito et al., 2012) or beta series correlations (Rissman et al., 2004), could yield both interesting differences and show areas of robustness in network profiles. However, this is well beyond the scope of the current project.

While our findings here provide strong evidence in support of the dynamical systems model, some caveats must be considered. First, all our analyses have been performed at the macroscopic-level. As mentioned above, even though evidence suggests that a similar behavior is expected at smaller network scales, future studies should test this *ex profeso*. On a related point, since the correlation matrices become computationally intractable at the voxel-level, and in order to maintain both activation and network measures with the same spatial resolution, we opted to perform all analyses at the region level, using a predefined parcellation template. This obviously introduces some degree of anatomically bounded spatial smoothing in the data, which may be contributing to inflating the similarities in both task-related activation and network profiles between tasks. Smoothing would be problematic if we were interested in null hypotheses tests on spatial patterns (Markello and Misic, 2021), however, the analysis used here does not rely on such spatial hypothesis testing. Thus, this region-level approach does not invalidate the main conclusions of our study that similarity in the topology of activation patterns does not perfectly associate with similarity in network architecture. Finally, one might question whether the BOLD time series first needed to be deconvolved with the hemodynamic response function prior to estimating the edge time series. It has been argued that deconvolution in block-design tasks, like our Stroop task and MSIT, may not be necessary (Di and Biswal, 2017; Di et al., 2020). However, it is important to point out that while changing the choices in the preprocessing and analysis steps may lead to nuanced differences in certain aspects of our results, none of these potential limitations would likely change the primary conclusion we have drawn from our observations.

Regardless of these limitations, our results clearly illustrate that important aspects of task representations are encoded in the associations between regions, which are unique to and complement information reflected in the spatial topology of activation (Gratton et al., 2016; Chan et al., 2017). Indeed, our findings bolster previous work looking at informational connectivity (Coutanche and Thompson-Schill, 2013), that highlights the information value of associations between regions in understanding task representations. Further work should dig deeper into the high dimensional relationships between localized activation and global connectivity dynamics when trying to understand the nature of representations int he brain.

## Code and data availability

The code used to generate all the analyses and results can be found in https://github.com/CoAxLab/cofluctuating-task-connectivity.

## Acknowledgements

Research reported in this publication was supported by the National Heart, Lung, snd Blood Institute of the National Institutes of Health under Award Numbers P01 HL040962 and R01 HL 1089850. The content is solely the responsibility of the authors and does not necessarily represent the official views of the National Institutes of Health.

